# Increased Ca^2+^ signaling through Ca_V_1.2 induces tendon hypertrophy with increased collagen fibrillogenesis and biomechanical properties

**DOI:** 10.1101/2023.01.24.525119

**Authors:** Haiyin Li, Antonion Korcari, David Ciufo, Christopher L. Mendias, Scott A. Rodeo, Mark R. Buckley, Alayna E. Loiselle, Geoffrey S. Pitt, Chike Cao

**Affiliations:** Center for Musculoskeletal Research, University of Rochester Medical Center, Rochester, NY; Department of Orthopeadics, University of Rochester Medical Center, Rochester, NY; Department of Biomedical Engineering, University of Rochester Medical Center, Rochester, NY; Arizona Bone, Joint and Sports Medicine Center, Phoenix, AZ; Sports Medicine and Shoulder Service, Hospital for Special Surgery, New York, NY; Cardiovascular Research Institute, Weill Cornell Medicine, New York, NY

## Abstract

Tendons are tension-bearing tissues transmitting force from muscle to bone for body movement. This mechanical loading is essential for tendon development, homeostasis, and healing after injury. While Ca^2+^ signaling has been studied extensively for its roles in mechanotransduction, regulating muscle, bone and cartilage development and homeostasis, knowledge about Ca^2+^ signaling and the source of Ca^2+^ signals in tendon fibroblast biology are largely unknown. Here, we investigated the function of Ca^2+^ signaling through Ca_V_1.2 voltage-gated Ca^2+^ channel in tendon formation. Using a reporter mouse, we found that Ca_V_1.2 is highly expressed in tendon during development and downregulated in adult homeostasis. To assess its function, we generated *ScxCre;Ca_V_1.2^TS^* mice that express a gain-of-function mutant Ca_V_1.2 channel (Ca_V_1.2^TS^) in tendon. We found that tendons in the mutant mice were approximately 2/3 larger and had more tendon fibroblasts, but the cell density of the mutant mice decreased by around 22%. TEM analyses demonstrated increased collagen fibrillogenesis in the hypertrophic tendon. Biomechanical testing revealed that the hypertrophic Achilles tendons display higher peak load and stiffness, with no changes in peak stress and elastic modulus. Proteomics analysis reveals no significant difference in the abundance of major extracellular matrix (ECM) type I and III collagens, but mutant mice had about 2-fold increase in other ECM proteins such as tenascin C, tenomodulin, periostin, type XIV and type VIII collagens, around 11-fold increase in the growth factor of TGF-β family myostatin, and significant elevation of matrix remodeling proteins including Mmp14, Mmp2 and cathepsin K. Taken together, these data highlight roles for increased Ca^2+^ signaling through Ca_V_1.2 on regulating expression of myostatin growth factor and ECM proteins for tendon collagen fibrillogenesis during tendon formation.

## Introduction

Tendons are tension-bearing tissues transmitting force from muscle to bone for body movement. This specialized function of tendons is supported by tendon specific extracellular matrix (ECM) structure with hierarchically organized collagen fibrils into fibers, fascicles, and then tendon (Silver, Freeman, & Seehra, 2003). Collagen fibrils are composed primarily of type I collagen, a triple helical polypeptide chains encoded by the genes *Col1a1* and *Col1a2*. Other tendon components are also present in the ECM important for tendon fibrillogenesis, such as type III, V, VI, XII and XIV collagens, small leucine rich proteoglycans (e.g., decorin, aggrecan, biglycan, and fibromodulin), and glycoproteins (e.g., tenascin C, tenomodulin, fibronectin, elastin, and collagen oligomeric matrix protein)(Mouw, Ou, & Weaver, 2014).

Within the organized collagen fibrils reside tendon cells which are constantly subjected to mechanical stimulation *in vivo*, including shear stress, tensile loading, and compressive force. Both tendon fibroblasts, the major cells in tendon, and the tendon stem or progenitor cells (TSPCs) sense and respond to a variety of mechanical loads (C. Zhang, Zhu, Zhou, Thampatty, & Wang, 2019; J. Zhang & Wang, 2013), which are then converted to cellular responses and biological signals, a process called mechanotransduction (Dunn & Olmedo, 2016; Ingber, 2006). Normal physiological loads are required for tendon fibroblast differentiation, ECM synthesis and organization during tendon development and adult tendon homeostasis (Bhole et al., 2009; Henderson & Carter, 2002; Kalson et al., 2011; Nowlan, Murphy, & Prendergast, 2007; Subramanian, Kanzaki, Galloway, & Schilling, 2018). In contrast, underloading or overloading results in decreased synthesis of tendon ECM proteins, ECM degeneration or aberrant TSPC differentiation (Wang, Guo, & Li, 2012). However, the molecular mechanisms of tendon mechanobiology remains unclear.

Ca^2+^, a ubiquitous intracellular signal, controls many cellular functions including muscle contraction, immune cell activation, gene transcription and cell proliferation (Clapham, 2007). A transient increase of intracellular Ca^2+^ concentration ([Ca^2+^]_i_) has been reported as one of the earliest responses upon mechanical stimulation, activating various physiological and pathological functions in tendon, ligaments, bone, cartilage, and muscle development, suggesting increased [Ca^2+^]_i_ plays a key role in mechanotransduction [reviewed in (Wall & Banes, 2005; Wall et al., 2016)]. In tendons, the Snedeker group recently showed with Ca^2+^ imaging that a transient [Ca^2+^]_i_ increase was observed by mechanical stimulation of the rat tail tendon fascicle *ex vivo* or isolated rat and human tenocytes; *Piezo1* loss-of-function, gain-of-function, and pharmacological approaches identified this stretch-activated ion channel as the mechano-sensor in tendon cell mechanotransduction, regulating tendon tissue stiffness (Passini et al., 2021). However, ion influx through Piezo1 is rapid, transient, and not specific to Ca^2+^ (Coste et al., 2010). Other source of Ca^2+^ entry may be required for the mechanical stimulated Ca^2+^ response in tendon cells. It could be functionally coupled with Piezo1 activation upon mechanical stimulation for opening and induces substantial Ca^2+^ signal for tendon cell mechanotransduction. For example, the Ca_V_1.2 channel has been shown to mediate mechanosensitive Ca^2+^ influx in intestinal smooth muscle cells (Lyford et al., 2002) and is broadly expressed (Pitt, Matsui, & Cao, 2021).

Ca_V_1.2 belongs to the family of L-type voltage-gated Ca^2+^ channels (L-VGCCs), mediating Ca^2+^ influx into the cell upon membrane depolarization. Ca_V_1.2 channel is composed of α_1C_, β and α2δ subunits (Catterall, 2000), among which the α_1C_ subunit is the ion conducting pore while the β and α2δ are auxiliary subunits that modulate channel properties (Serysheva, Ludtke, Baker, Chiu, & Hamilton, 2002). Ca_V_1.2 is highly voltage-dependent, which is characterized by an activation threshold at a membrane potential around −20 mV (Catterall, Perez-Reyes, Snutch, & Striessnig, 2005). Most studies on Ca_V_1.2 focused on its function in excitable cells such as cardiomyocytes and neurons, where action potentials facilitate activation of this voltage-gated channel. However, few studies focused on Ca_V_1.2 in non-excitable cells, which have more restricted changes in membrane potential. Interestingly, studies of Timothy syndrome (TS), a multiorgan disorder (e.g., cardiac arrhythmias, autism, syndactyly and craniofacial abnormalities), caused by a *de novo* G406R mutation in the Ca_V_1.2 pore forming α1C subunit encoded by *CACNA1C* (Splawski et al., 2005; Splawski et al., 2004), revealed critical but previously unappreciated roles for Ca_V_1.2 in many non-excitable cells. For example, digital and craniofacial abnormalities in TS patients suggest roles for Ca_V_1.2 in development and morphogenesis, and that aberrant G406R mutant channel (Ca_V_1.2^TS^) activity adversely affects canonical developmental signals. Consistent with these hypotheses, we observed robust Ca_V_1.2 endogenous expression in osteoblast progenitors during craniofacial and limb development using a Ca_V_1.2 lacZ reporter mouse line (C. Cao et al., 2017; Kapil V. Ramachandran et al., 2013). In addition, by driving a *Ca_V_1.2^TS^* transgene with *Prx1Cre, Col1a1Cre, Col2aCre* or *Sp7Cre*, we demonstrated that Ca_V_1.2^TS^ mutant channels promote bone formation via increased osteoblast differentiation and decreased osteoclast function. This also prevented estrogen deficiency-induced bone loss, highlighting the unexpected role for Ca_V_1.2 in non-excitable tissue (Cao et al., 2019; C. Cao et al., 2017; Kapil V. Ramachandran et al., 2013). The G406R mutation impairs Ca_V_1.2 channel inactivation (closing) leading to more Ca^2+^ ions to flow into the cytoplasm. Thus, the consequences of *Ca_V_1.2^TS^* mutant channel expression reported above result from a gain-of-function effect. However, whether Ca_V_1.2 confers analogous effects during tendon development is not known. To address this, we tested whether the L-VGCC Ca_V_1.2 is expressed in non-excitable tendon tissue, and whether an increase of Ca^2+^ signaling through *Ca_V_1.2^TS^* mutant channels in tendon affects tendon formation.

## Results

### Ca_V_1.2 is expressed in tendon fibroblasts during mouse tendon development and early postnatal growth

To determine whether the Cav1.2 channel contributes to tendon formation during development, postnatal growth and homeostasis, we first elucidated Ca_V_1.2 channel expression in tendons at different stages by using a Ca_V_1.2 reporter mouse line (*Ca_V_1.2^+/lacZ^*, B6.129P2-*Cacna1c^tm1Dgen/J^*) in which the bacterial *lacZ* gene encoding β-galactosidase fused to a nuclear localization signal was knocked into *Cacna1c* labeling nuclei of cells that express Ca_V_1.2 (C. Cao et al., 2017; Kapil V. Ramachandran et al., 2013). We performed whole mount X-gal staining of the forelimbs and hindlimbs, followed by frozen sectioning and histological analysis. We found substantial X-gal staining in the developing digital tendons from E13.5 (Fig. 1A), and increased intensity of staining at late embryonic stages (Fig. 1B). X-gal staining in digital tendons was confirmed by histological analysis on frozen sections of the developing digits (Fig. 1C). Furthermore, we found *Ca_V_1.2* expression persisted through early postnatal stages (~P3) in the Achilles tendons (Fig. 1D) and patellar tendons (Fig. 1F). Notably, X-gal staining was exclusively localized in the nucleus, which is distinct from any non-specifical staining resulting from endogenous β-galactosidase activity, and thus provides an accurate picture of endogenous *Ca_V_1.2* expression. However, in adult tendons, *Ca_V_1.2* expression was dramatically downregulated. For example, adult Achilles tendon demonstrated restricted *Ca_V_1.2* expression mostly seen in the myotendinous junction with very sparse expression in the tendon substance (Fig. 1E). In contrast, adult patellar tendon retains strong *Ca_V_1.2* expression throughout the tendon tissue, but in relatively fewer cells compared with patellar tendons in early postnatal stage (Fig. 1G). In summary, the dynamic expression of *Ca_V_1.2* during tendon development, postnatal growth and adult homeostasis stage suggests that Ca_V_1.2 and its mediated Ca^2+^ signaling may play a critical role for tendon formation.

**Figure 1.**
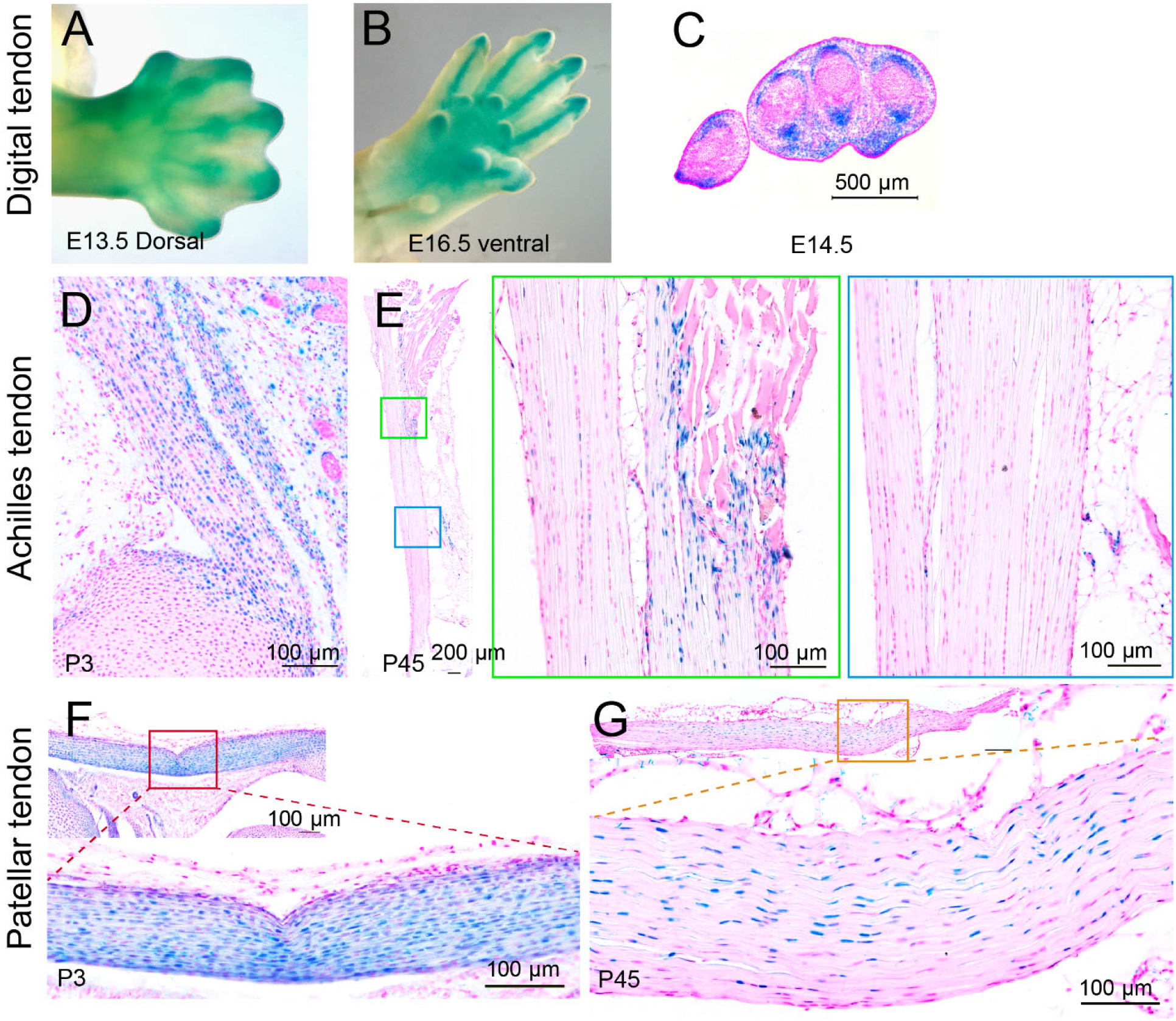
Ca_V_1.2 is expressed in tendon fibroblasts during embryonic development and early postnatal growth in mice. (A&B) Representative images of whole-mount X-gal stained forelimbs from *Ca_V_1.2^+/LacZ^* embryos harvested at E13.5 (A) and at E16.5 (B) are shown to illustrate expression of the transgene in the skeletal elements. (C-G) Fast red counterstaining was performed on frozen sections of whole-mount X-gal stained *Ca_V_1.2^+/LacZ^* forelimb digits at E14.5 (C), Achilles tendon at P3 (D) and P45 (E), and patellar tendon P3 (F) and P45 (G); boxed regions were obtained at high power. The nuclear localization of blue color resulting from X-gal staining illustrates the specific transgene expression in tendon fibroblasts.

### Expression of *Ca_V_1.2^TS^* mutant channel in *ScxCre^+^* lineage cells enhances tendon formation

To investigate the role of Ca_V_1.2 on tendon formation *in vivo*, we exploited the conditional transgenic mouse line carrying the *Ca_V_1.2^TS^* mutant cDNA in the *Rosa26* locus (S. P. Paşca et al., 2011). We generated *ScxCre;Ca_V_1.2^TS^* mice to allow for *Ca_V_1.2^TS^* expression under the *Scleraxis (Scx)* promoter, which regulates transcription during tendon development, and is the earliest known marker of tendon progenitors (Schweitzer et al., 2001). Macroscopic observation of *ScxCre;Ca_V_1.2^TS^* mutant mice revealed hypertrophic tendons in all tendons examined, including Achilles tendon, plantaris tendons, patellar tendons, tail tendons, tendons in the forelimbs and back tendons, compared to tendons in control mice at one month of age (Fig. 2). Histologic analyses further confirmed tendon hypertrophy in *ScxCre;Ca_V_1.2^TS^* mutant mice (Fig. 3). Fast green and hematoxylin staining revealed that the cross-section area (CSA) was increased by 61% in patellar tendons, 70% in plantaris tendons, and 74% in Achilles tendons from 1-month-old *ScxCre;Ca_V_1.2^TS^* mutant mice compared with those in control mice. Consistently, cell numbers in patellar tendons, plantaris tendons and Achilles tendons were increased by 21%, 32% and 38% in the mutant mice, respectively, indicating increased cell proliferation in the mutant tendons. However, *ScxCre;Ca_V_1.2^TS^* mutant mice had decreased cell density (cell number/CSA) by 22% in all three types of tendons. This suggests that *Ca_V_1.2^TS^*-expressing tendon fibroblasts are more functionally active. Moreover, we found that *ScxCre;Ca_V_1.2^TS^* mice had thicker tail tendon fascicles, which have a broader size distribution than those in the control mice (Supplementary Fig. 1). Notably, there was no change in the number of fascicles of both ventral and dorsal tail tendons between genotypes, suggesting that Ca_V_1.2^TS^ mutant channels affect tendon fascicle growth but not fascicle determination.

**Figure 2.**
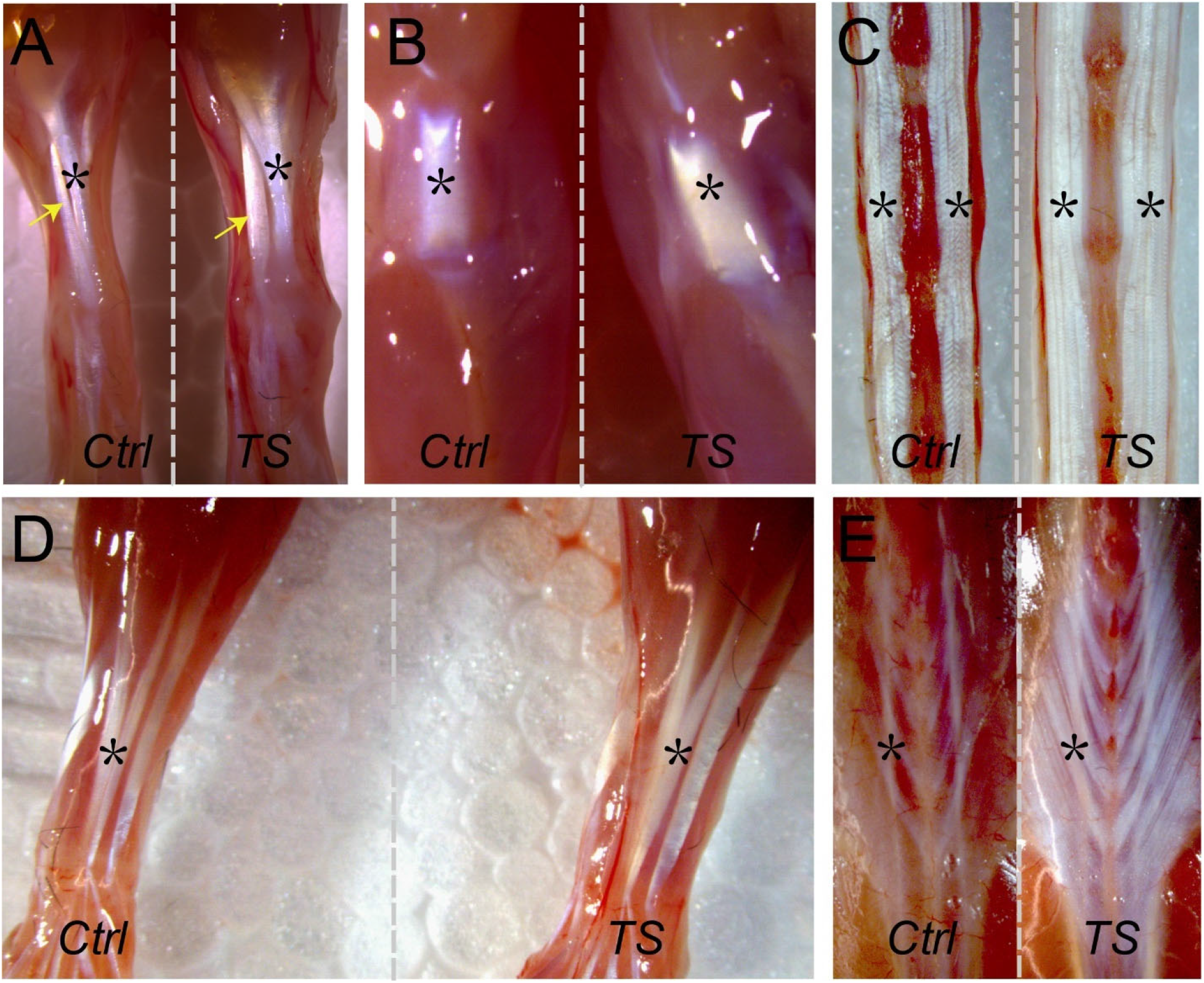
Enhanced tenogenesis in *ScxCre; Ca_V_1.2^TS^* mice. Gross anatomy was performed on tendons from 1 month-old *Cre^-^; Ca_V_1.2^TS^* and *ScxCre;Ca_V_1.2^TS^* mice, and representative images of: plantaris tendons (arrows) and Achilles tendons (*) in hindlimbs (A), patellar tendons (B), tail tendons (C), tendons in forelimbs (D) and back tendons (E) are shown. Note the white tissues are tendons indicated by * or arrows, and appear larger in *ScxCre;Ca_V_1.2^TS^ (TS)* vs. *Cre^-^; Ca_V_1.2^TS^* control (*Ctrl*) mice.

**Figure 3.**
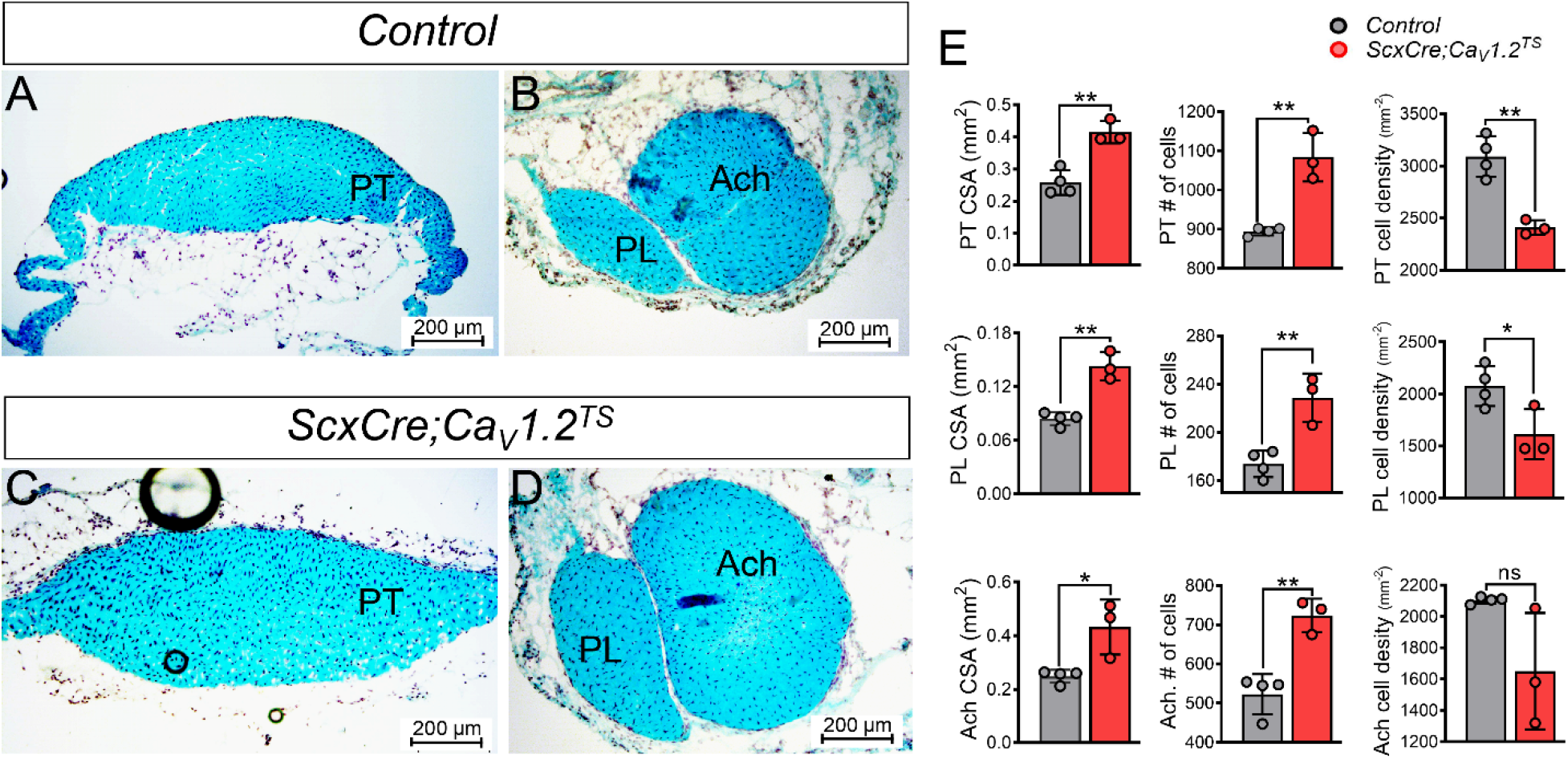
Increased tendon hypertrophy in *ScxCre;Ca_V_1.2^TS^* mice. (A, B, C and D) Representative images of fast green and hematoxylin stained tendon cross sections from littermate control (*Cre^-^;Ca_V_1.2^TS^*) and *ScxCre; Ca_V_1.2^TS^* mice at 1 month of age. (E) Histomorphometry was performed on these sections to quantify tendon cross section area (CSA), number of cells in whole tendon cross section area, and tendon cell density (number of cells/CSA), and the data for each tendon are presented with the mean for the group ± SD (n >= 3, * *p* < 0.05, ** *p* < 0.01, ns, not significant). Statistical analysis was performed by 2-tailed unpaired *t* test, and a *p* value less than 0.05 was considered significant. PT: patellar tendon, PL: plantaris tendon, Ach: Achilles tendon.

### Expression of Ca_V_1.2^TS^ mutant channel in *ScxCre^+^* lineage cells alters tendon collagen fibril size distribution

To further investigate the effects of the Ca_V_1.2^TS^ mutant channel on tendon collagen fibrillogenesis, we performed ultrastructural analyses using transmission electron microscopy (TEM) of Achilles tendons from *ScxCre;Ca_V_1.2^TS^* mutant mice and littermate controls at 1 month of age. In the mutant Achilles tendons, the collagen fibrils displayed normal circular cross-sectional profiles, similar to the control fibrils (Fig. 4A and B). However, the collagen fibril density in the mutant tendons was increased by 68% without a significant change in interfibrillar spacing (Fig. 4C and D), indicating increased collagen fibrillogenesis in *ScxCre;Ca_V_1.2^TS^* mutant mice. Furthermore, the mutant Achilles tendons were packed with more small- to-middle size collagen fibrils, resulting in the change of the repartition of collagen fibrils in *ScxCre;Ca_V_1.2^TS^* mutant mice (Fig. 4E). In addition, a leftward shift of the fibril size distribution of the mutant tendon in the cumulative fraction analysis further supported that the collagen fibrils in mutant mice are smaller than those in the control mice (p <0.01 by Kolmogorov-Smirnov test) (Fig. 4F). Taken together, these data suggest increased Ca^2+^ signaling through the Ca_V_1.2^TS^ mutant channel increases collagen fibril assembly, which contributes to tendon hypertrophy in *ScxCre;Ca_V_1.2^TS^* mutant mice.

**Figure 4.**
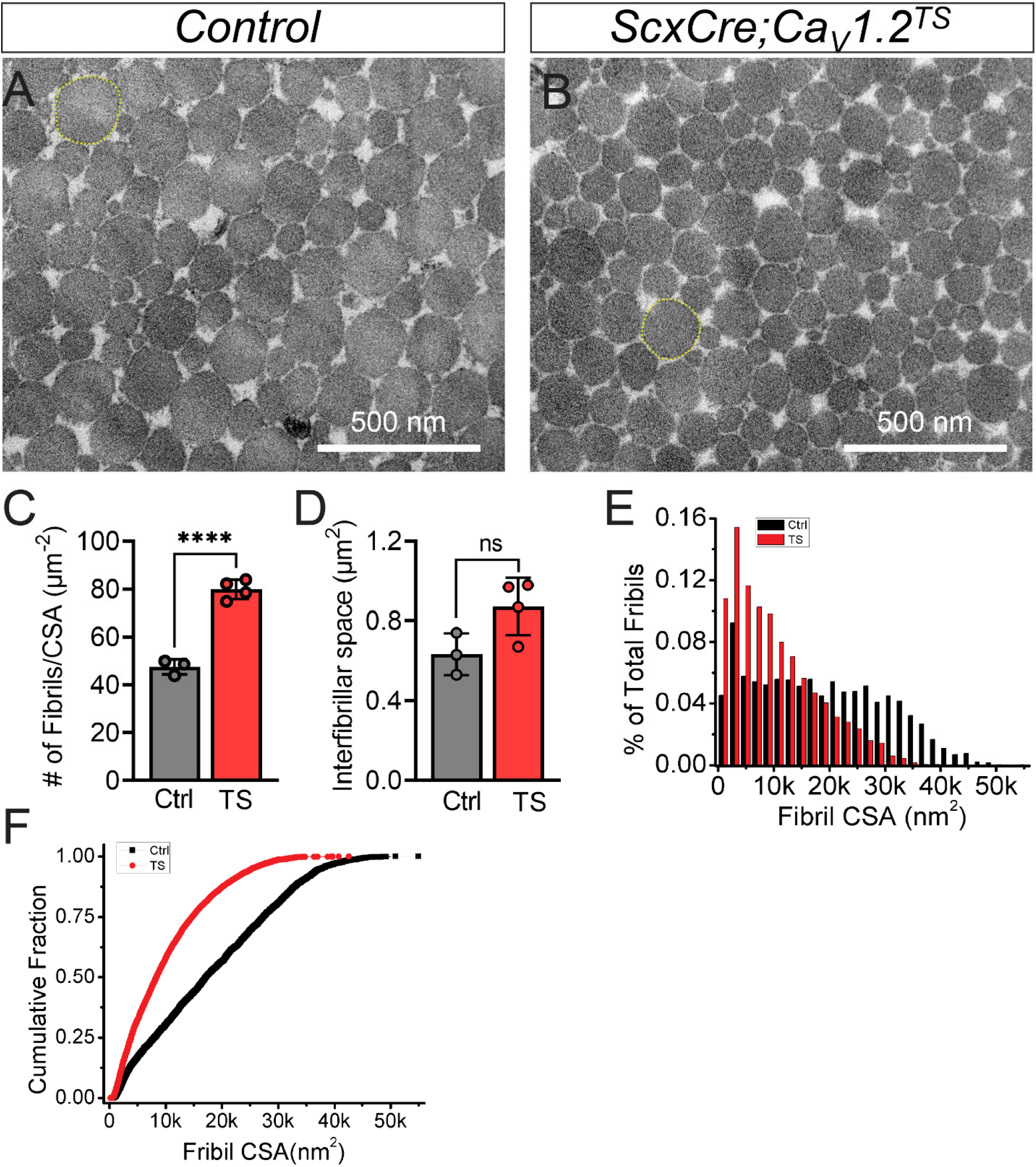
Altered collagen fibril size distribution of Achilles tendons in *ScxCre;Ca_V_1.2^TS^* mice. (A and B) Representative transmission electron microscopy (TEM) images of Achilles tendon collagen fibrils from littermate control (*Cre^-^;Ca_V_1.2^TS^*) and *ScxCre;Ca_V_1.2^TS^* mice at 1 month of age are shown to illustrate the smaller fibrils in *ScxCre;Ca_V_1.2^TS^* mice as illustrated by the difference in size of the largest fibril in each group (yellow highlight). (C and D) Histomorphometry was performed on these TEM sections to quantify fibril density (number of fibrils/CSA of fibrils) (C) and the collagen interfibrillar spacing (D); the data are presented for each tendon with the mean ± SD (n >= 3, \*\*\*\**p* < 0.0001). (D) Histograms showing the altered frequencies of collagen fibril CSAs from *ScxCre;Ca_V_1.2^TS^* mice compared to control mice. (E) Cumulative fraction analysis of collagen fibril CSAs, showing the left-shift of the size of collagen fibrils in *ScxCre;Ca_V_1.2^TS^* mice. Kolmogorov-Smirnov test shows that the tendon fibrils are significantly smaller than those in control mice, n >= 3, *p* < 0.001. CSA: cross section area. Ctrl: *Cre^-^;Ca_V_1.2^TS^* control mice; TS: *ScxCre;Ca_V_1.2^TS^* mice.

### Ca_V_1.2^TS^ alters tendon biomechanical properties

Since cellular arrangement and fibril packing are important determinants of biomechanical properties (Heather L. Ansorge et al., 2009; Dunkman et al., 2013; Thorpe, Udeze, Birch, Clegg, & Screen, 2012), increased tendon growth and the change of collagen fibril size distribution in *ScxCre;Ca_V_1.2^TS^* mutant tendons may alter their mechanical properties. Therefore, we performed the biomechanical property test in mature mutant Achilles tendon in comparison with the control ones. Cross-sectional area (CSA), peak load, peak stress, stiffness, and elastic modulus were measured. Mutant Achilles tendons displayed a significantly larger CSA than the control Achilles tendons (Fig. 5A), exhibited ~1.50-fold increase in peak load, and a ~1.52-fold increase in stiffness (tensile/displacement) in the force-displacement response (Fig. 5B and C), indicating functional gain in structure properties of mutant Achilles tendons. However, the material properties including the peak stress (the peak load per unit area), and the elastic modulus (a measurement of the stiffness of an isotropic elastic material per unit area), did not show significant changes between genotypes (Fig. 5D and E). This suggests that the increase in structural stiffness in *ScxCre;Ca_V_1.2^TS^* mutant tendons is due to the increased tendon mass.

**Figure 5.**
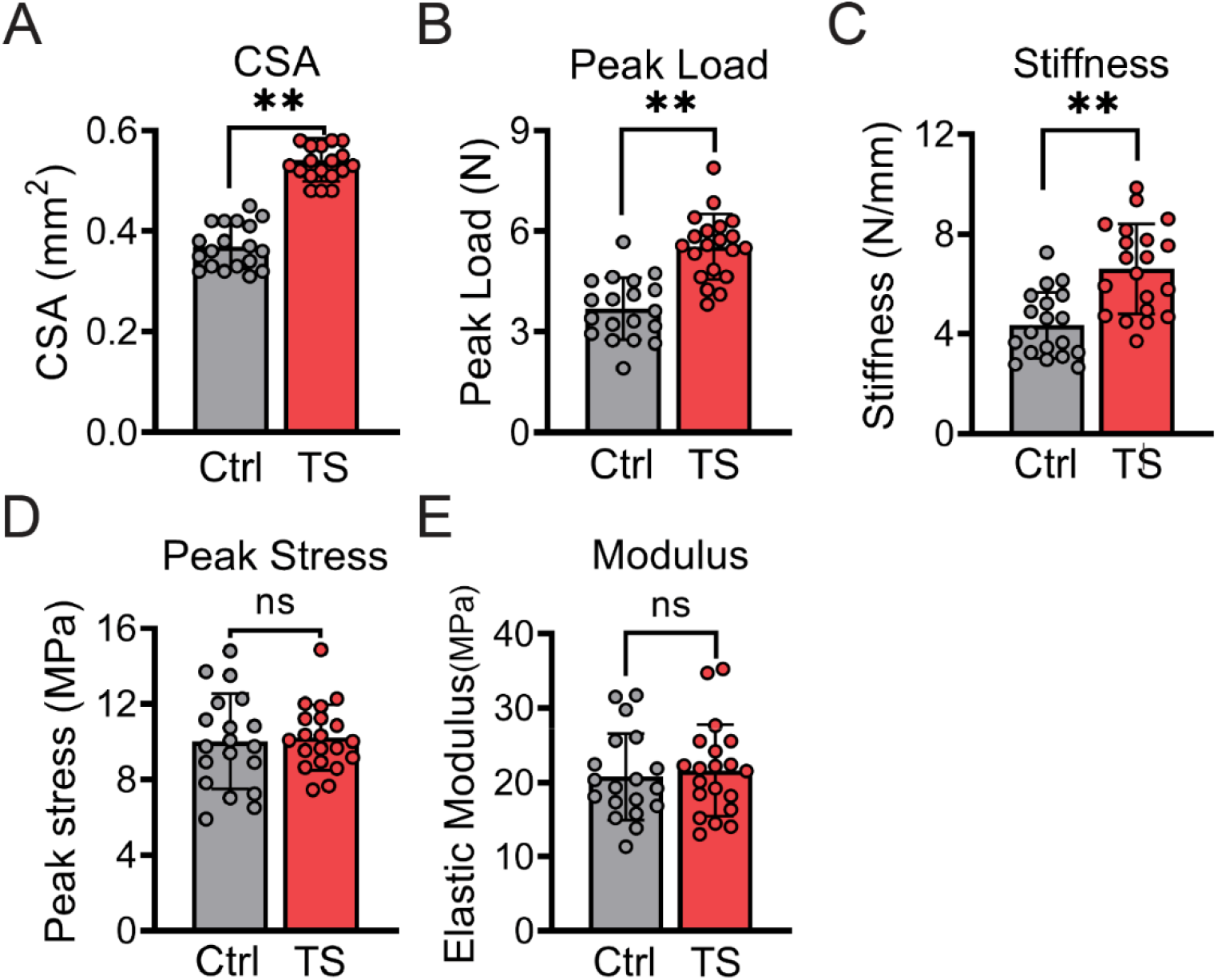
Specific biomechanical alteration of Achilles tendons in *ScxCre;Ca_V_1.2^TS^* mice. Uniaxial displacement-controlled stretching test was performed on Achilles tendons from control and *ScxCre;Ca_V_1.2^TS^* of both male and female mice at 1 month of age to quantify: peak load (A), stiffness (B), elastic modulus (D) and peak stress (E) mice. The data are presented for each tendon with the mean ± SD (n >= 19, ** *p* < 0.01, ns: not significant via *t*-test).

### Ca_V_1.2^TS^ alterations in the proteome of tendons

To define the molecular consequences of Ca_V_1.2^TS^ expression, we quantified tissue-wide protein changes using mass spectrometry-based proteomic analysis on *ScxCre;Ca_V_1.2^TS^* versus control mice at 1 month of age. We observed 89 upregulated proteins (>1.5 fold-change) and 102 downregulated proteins (<-1.5 fold-change) in Ca_V_1.2^TS^-expressing mice compared with those in control mice (Fig. 6A and B, Supplementary Fig. 2). The six proteins identified with the largest increase in expression in *ScxCre;Ca_V_1.2^TS^* mutant tendons were Mstn (myostatin, a member of the TGFβ-superfamily), Pavlb (a high affinity Ca^2+^-binding protein similar to calmodulin in structure and function), Tnn (Tenascin-N), Cthrc1 (collagen triple helix repeat-containing protein 1), Angptl1 and Antptl2 (angiopoietin-like proteins). In contrast, the proteins with the largest decrease were Chad (chondroadherin, a cartilage matrix protein), Zmym4 (Zinc Finger MYM-type containing 4), Angptl7, Htra4 (high temperature requirement factor A4) and Omd (Osteomodulin) (Fig. 6C). Moreover, Gene Ontology (GO) enrichment analyses were performed to classify the putative functions of the differentially upregulated and downregulated protein sets in *ScxCre;Ca_V_1.2^TS^* mice in comparison with those of the control mice. In GO terms of cellular component, we found that these differentially expressed proteins are related to ECM, Proteinaceous ECM, extracellular region, extracellular exosome, and extracellular space (Fig. 6D). Furthermore, GO analysis in term of biological process showed that many of these proteins were involved in ECM organization (increase of Col8a1, Mmp14, Mmp2, and Postn, decrease of Abi3bp, Ccdc80, Col15a1, Col24a1, Fbln1, Fbln2, Lgals3, Prdx4, Tnxb, Vit and Vtn), collagen fibril organization (increase of Col14a1, decrease of Comp, Fmod, and Tnxb), collagen catabolic process (increase of Ctsk, Mmp14, and Mmp2), response to mechanical stimulus (increase of Mmp14, Mmp2, Postn, Tnc, and decrease of Thbs1 and Dcn) as shown in Fig. 6E. We didn’t observe significant differences in the abundance of type I collagen and type III collagen between genotypes.

**Figure 6.**
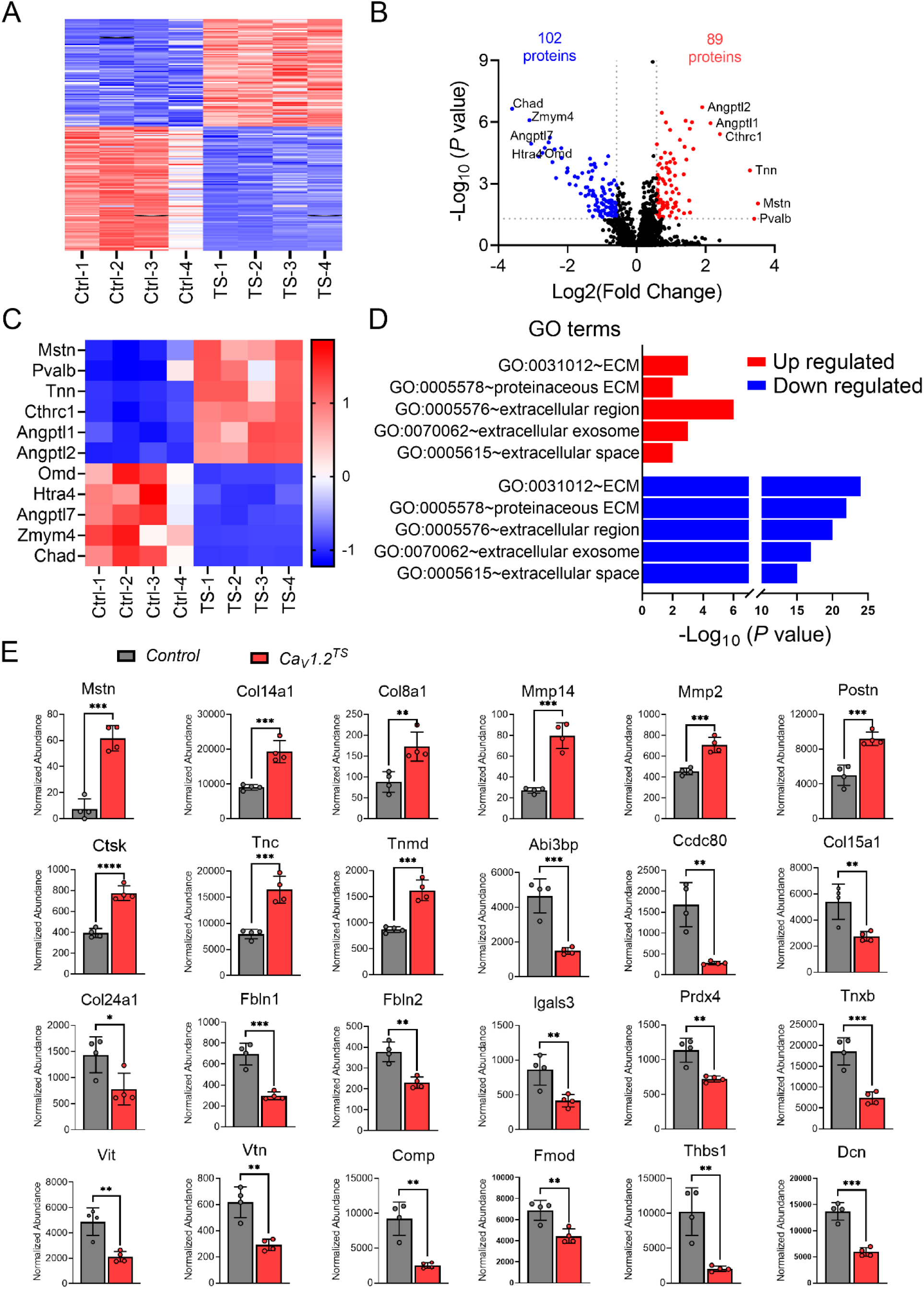
Altered proteomics of Achilles tendons in *ScxCre;Ca_V_1.2^TS^* mice. Proteomic analysis was performed on Achilles tendon from control (*Cre^-^;Ca_V_1.2^TS^*) and *ScxCre;Ca_V_1.2^TS^* mice at 1 month old. (A) Heatmap of all significantly different proteins between control and *ScxCre;Ca_V_1.2^TS^* mice. (B) Volcano plot showing upregulated (red) and downregulated (blue) protein expression in *ScxCre;Ca_V_1.2^TS^* mice compared to control mice. (C) Heatmap showing the proteins with largest increase and decrease in expression in *ScxCre;Ca_V_1.2^TS^* mice compared to control mice. (D) The GO terms of upregulated and downregulated differentially expressed proteins in *ScxCre;Ca_V_1.2^TS^* mice with DAVID analysis. (E) Normalized abundance of the proteins with significant increase and decrease in expression in *ScxCre;Ca_V_1.2^TS^* mice compared to control mice. Values are mean ± SD. * *p* < 0.05, ** *p* < 0.01, *** *p* < 0.001. n = 4 for each group. Identification of proteins, > 1.5-fold or <-1.5-fold change in abundance and FDR *p*-value <0.05 was considered significant.

To validate these findings, we performed real-time quantitative PCR (RT-qPCR) of selected markers related to tendon formation including: *Mstn, Tnc, Tnmd, Mmp14, Scx* and *Col1a1*, and found these gene mRNA expression was consistent with their expression at protein level (Fig. 7). For example, the growth factor Mstn, which is a positive regulator for tendon formation (Christopher L. Mendias, Konstantin I. Bakhurin, & John A. Faulkner, 2008) and had the greatest protein increase (~11.4-fold) in *ScxCre;Ca_V_1.2^TS^* tendons, displayed a ~35.7-fold upregulation in mRNA expression (Fig. 7A). The expression of *Tnc*, *Tnmd* and *Mmp14* was increased around 2.9-, 2.5- and 4.9-fold, irrespectively, in *Ca_V_1.2^G406R^*-expressing Achilles(Fig. 7B-D). In contrast, *Scx* and *Col1a1* didn’t show significant differences between genotypes at mRNA level (Fig. 7E and F). Taken together, these data suggest that Ca_V_1.2^TS^ mutant channels promoted tendon formation by upregulating expression of Mstn and the less abundant ECM proteins for tendon collagen fibril organization and ECM turnover.

**Figure 7.**
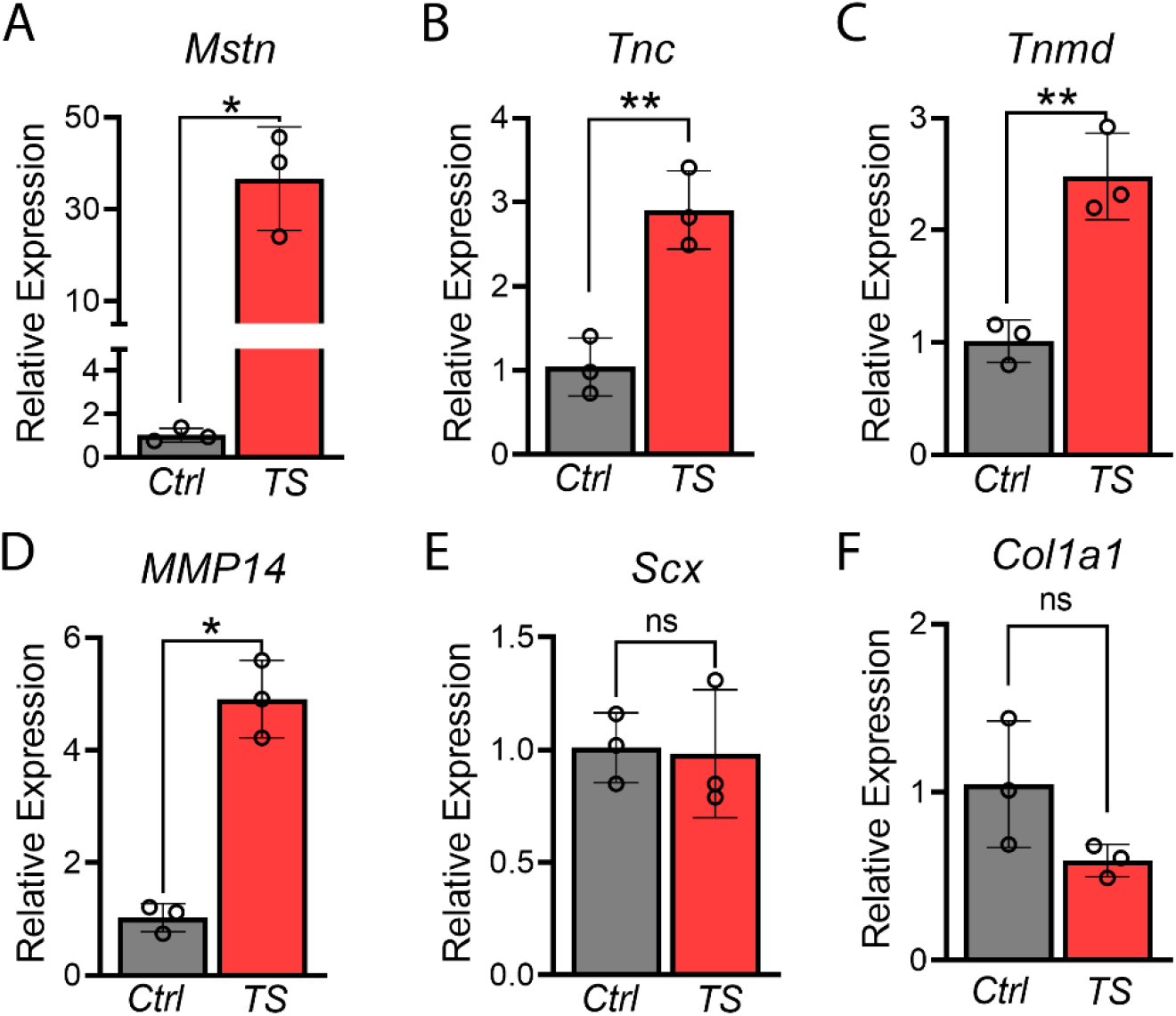
Gene expression analysis in Achilles/plantaris tendons of control and *ScxCre;Ca_V_1.2^TS^* mice. (A-F) Quantitative analysis of RT-qPCR for *Mstn, Tnc, Tnmd, Mmp14, Scx*, and *Col1a1* expression. Target gene expression values were normalized to the stable housekeeping gene *Gapdh*, and then to relative control expression levels. Values are mean ± SD, n = 3 for each group. Difference between groups were tested using 2-tailed unpaired *t* test (* *p* < 0.05, ** *p* < 0.01).

## Discussion

The current study identified the dynamic expression of endogenous *Ca_V_1.2*, a voltage-dependent Ca^2+^ channel in tendon fibroblasts during tendon development, growth and homeostasis by utilizing a conclusive Ca_V_1.2 lacZ reporter mouse line. This discovery prompted us to examine whether this voltage-dependent Ca^2+^ channel functions in tendon formation. Using a transgenic mouse model (*ScxCre;Ca_V_1.2^TS^*), we demonstrated that expression of the gain-of-function G406R mutant channel Cav1.2^TS^ specifically in *ScxCre^+^* tendon fibroblasts dramatically promotes tendon formation. Biomechanical testing showed that the enlarged tendons in *ScxCre;Ca_V_1.2^TS^* mutant mice display a dramatic increase of their structural properties including stiffness and peak load, but have similar material properties, such as peak stress and elastic modulus, compared to WT tendons. Therefore, the increased tendon stiffness and peak load could be owing to the proportional increase of tendon thickness (measured by CSA). Notably, these changes of tendon biomechanical properties in *ScxCre;Ca_V_1.2^TS^* mutant mice are comparable to those in the adaptation of human tendons after years of long-term training with larger tendon CSA and increased tendon stiffness, but no differences in material properties based on the meta-analysis of tendon property changes with training (Wiesinger, Kosters, Muller, & Seynnes, 2015), suggesting the role of Ca^2+^ signaling via Ca_V_1.2 may be linked with tendon loading and mechanotransduction. Taken together, our data provides the first evidence that modulating Ca^2+^ signaling through Ca_V_1.2 in tendon fibroblasts *in vivo* affects tendon formation, highlighting additional unexpected roles of Ca_V_1.2 channels in nonexcitable tissues that we previously reported (Cao et al., 2019; C. Cao et al., 2017; Kapil V. Ramachandran et al., 2013).

The tendon hypertrophy in *ScxCre;Ca_V_1.2^TS^* mutant mice is likely due to both increased tendon fibroblast proliferation and ECM collagen fibril formation. This speculation is supported by our findings that the mutant tendons had significantly more cells but reduced cellular density (Fig.3). This specific spatial organization of tendon cells can result from more increased collagen fibrillogenesis and fibril assembly relative to the increase in tendon cell proliferation. This is supported by our TEM findings that *ScxCre;Ca_V_1.2^TS^* mutant tendons generated more collagen fibrils with smaller diameter versus WT tendons (Fig. 4). During tendon development, short and small-diameter fibril intermediates are initially assembled, which serve as the building blocks to form large and long collagen fibrils in late stage of tendon growth (Nurminskaya & Birk, 1998). The accumulated small-to-medium size fibrils in the mutant tendons indicated an increased collagen fibrillogenesis and active fibril formation in response to upregulated Ca^2+^ signaling in *ScxCre;Ca_V_1.2^TS^* mice. Subsequently, we performed proteomic analysis to define the tissue-wide protein change and understand the molecular mechanisms responsible for the altered collagen fibrillogenesis. It is known that tendon collagen fibrillogenesis can be regulated in several ways, including 1) the synthesis of fibril-forming collagen (predominantly type I collagen with varying amounts of type II, III and V in tendon), 2) the abundance of specific fibril-associated proteoglycans and glycoproteins (such as decorin, biglycan, lumican, fibromodulin and COMP), or 3) the abundance of the fibril-associated collagen with interrupted triple helics (FACIT) (such as type IX, XII, XIV, XVI, XIX, XX, XXI, XXII, and XXVI collagens) (Kadler, Baldock, Bella, & Boot-Handford, 2007; Mouw et al., 2014; Nurminskaya & Birk, 1998). A deficiency of decorin, lumican, fibromodulin, COMP and type XVI collagen *in vivo* has been shown to result in larger and disorganized collagen fibrils (H. L. Ansorge et al., 2009; Chakravarti et al., 1998; Danielson et al., 1997; Piróg et al., 2010; Svensson et al., 1999). Interesting, our proteomic analysis showed that in *ScxCre;Ca_V_1.2^TS^* mutant tendons, the major fibril-forming collagens (Col1a1 and Col1a2) was not differentially expressed versus control tendons. However, the fibril-associated FACIT collagen (Col14a1) was significantly upregulated, while proteoglycans including decorin and fibromodulin and glycoprotein COMP were downregulated in *ScxCre;Ca_V_1.2^TS^* tendons. Thus, while a combination of these fibril-associated macromolecules with variable amount contributed to the tendon collagen fibrillogenesis and growth, type XIV collagen may be the dominant factor affecting collagen fibrillogenesis and assembly in *ScxCre;Ca_V_1.2^TS^* tendons. Furthermore, collagen fibrillogenesis requires activities of matrix metalloproteinases (MMPs) and their corresponding tissue inhibitors (TIMPs) to convert procollagen into collagen by removing the N- and C-pro-peptides, and alter the surface of fibril intermediates and/or interfibrillar matrix (Jones et al., 2006; Mouw et al., 2014). Notably, we found that Mmp2 and Mmp14 were both dramatically upregulated while Timp3 was downregulated in *ScxCre;Ca_V_1.2^TS^* tendons by proteomics analysis (Fig. 6E and 6G). It is known that Mmp2 is initially produced as latent pro-Mmp2, which requires the membrane type (MT) MMPs such as Mmp14 for cleavage and activation (Deryugina et al., 2001; Strongin et al., 1995). It has been shown that in an *in vitro* system, knockdown of Mmp14 inhibited proMmp2 activation (Wilkinson et al., 2012). Moreover, Mmp14 has been shown to promote new formation of collagen fibers, the high order of tendon structure; the tendons of mutant mice lacking Mmp14 have fewer collagen fibers than normal mice (Taylor et al., 2015). Whereas all active MMPs can be inhibited by TIMPs, Timp3 is the main TIMP which inhibits activity against some of the ADAMS and ADAMTS metalloproteinases (Del Buono, Oliva, Osti, & Maffulli, 2013; Mochizuki & Okada, 2007). Taken together, decrease in Timp3 expression along with the elevated expression of Mmp2, Mmp14 and Type XIV FACIT collagen in *ScxCre;Ca_V_1.2^TS^* mutant tendon will facilitate procollagen maturation, collagen fibril assembly and remodeling, all of which contribute to the active collagen fibrillogenesis and the higher structure fiber growth, ultimately tendon hypertrophy upon increased Ca^2+^ signaling.

Our proteomic analysis also identified a dramatic increase of myostatin in *ScxCre;Ca_V_1.2^TS^* mutant tendons. Myostatin, also called growth/differentiation factor-8 (GDF-8), is the growth factor of the transforming growth factor-β (TGF-β) superfamily. Myostatin is mostly known as a negative regulator of muscle growth, and the loss of myostatin function is associated with hypermuscular phenotypes in mice and cattle (Alexandra C. McPherron, Lawler, & Lee, 1997; A. C. McPherron & Lee, 1997). In contrast, myostatin was found to be a positive regulator for tendon formation as myostatin-deficient mice have small (a decrease in fibroblast number) and brittle tendons (a higher peak stress, a lower peak strain and increased stiffness) (C. L. Mendias, K. I. Bakhurin, & J. A. Faulkner, 2008). Thus, upregulation of myostatin in *ScxCre;Ca_V_1.2^TS^* mutant tendon may promote tendon hypertrophy in the mutant mice upon increased Ca^2+^ signaling. However, whether myostatin signaling mediates tendon growth in *ScxCre;Ca_V_1.2^TS^* mice requires further investigation. Conditional knockout of *Mstn* alleles in Ca_V_1.2^TS^-expressing tendon fibroblasts would be necessary to exclude the possibility that other myostatin-independent signaling pathways may also contribute to Ca_V_1.2^TS^-induced tendon formation. Furthermore, previous studies have shown that the p38 mitogen-activate protein kinase (MAPK) and Smad2/3 signaling cascades (Lee & McPherron, 2001; Philip, Lu, & Gao, 2005) in tendon fibroblasts were activated in response to myostatin treatment, which are required for the increased cell proliferation and gene expression including *Scx*, *Col1a1*, and *Tnmd* (C. L. Mendias et al., 2008). Consistently, we observed increased expression of *Tnmd* both at protein and mRNA levels in *ScxCre;Ca_V_1.2^TS^* mice. However, we didn’t detect a significant change of *Scx* and *Col1a1* expression in response to upregulated myostatin in *Ca_V_1.2^TS^*-expressing tendons. This discrepancy may be due to the maximal dose of myostatin normally used in *in vitro* cell culture while in *in vivo* system, myostatin may be dominantly maintained in an inactive form. Nevertheless, upregulation of *Tnmd* and *Tnc*, known as the downstream targets of transcription factor Scx and regulated by pSmad2/3 pathway (Berthet et al., 2013; Shukunami, Takimoto, Oro, & Hiraki, 2006), didn’t depend on a corresponding increase of *Scx* expression in *ScxCre;Ca_V_1.2^TS^* tendons.

Although the mechanism by which Ca_V_1.2 activation occurs in non-excitable tendon fibroblasts has yet to be explored, our finding that Ca_V_1.2 is dynamically expressed during tendon development, growth and adult homeostasis suggests a role of Ca^2+^ signaling in tendon fibroblasts spatiotemporally. It also implies the potential mechanisms to activate the voltage-gated Ca^2+^ channel. In adults, Ca_V_1.2 expression is dramatically decreased compared to that during tendon development and early postnatal growth. Ca_V_1.2 expression in adult Achilles tendon is restricted to the myotendinous junction (Fig. 1), a site where forces generated by myofibrils are transmitted across the cell membrane to act on tendon (Tidball & Lin, 1989). Notably, at this interface between an excitable muscle and non-excitable tendon, topological action potential may occur via gap junction (Ori et al., 2022), which results in non-excitable tendon fibroblasts in myotendinous junctions electrically excitable and then activate this voltage-dependent Ca^2+^ channel. Tendon fibroblasts in myotendinous junction could function as a signal initiator, which once stimulated by muscle contraction, will diffuse signal factors down to the neighboring fibroblasts. However, during stages before the stable myotendinous junction forms in tendons, usually at 1 month old in rats (Curzi, Ambrogini, Falcieri, & Burattini, 2013), or in patellar tendons (ligaments precisely) without the myotendinous junction structure, other mechanisms to activate Ca_V_1.2 voltage-dependent Ca^2+^ channel may also exist, by which the depolarizing drive does not require action potentials. Recently, the mechanosensitive channel Piezo1 has been reported to be expressed in tendon tissue and to sense the mechanical loading in tenocytes; knocking out Piezo1 in cultured tenocytes greatly decreases shear stress-induced Ca^2+^ signals (Passini et al., 2021). Given the fact that Piezo1 and its homolog Piezo2 conduct a rapid and transient cation influx (non-selective to Ca^2+^) (Coste et al., 2010), activation of Piezo1/2 by mechanical stimuli may provide an inward depolarizing current, which in turn activates the voltage-gated Ca_V_1.2 channels co-expressed in tendon fibroblasts to amplify the Ca^2+^ signal. However, whether Ca_V_1.2 is required for mechanotransduction in tendon has yet to be determined. Nevertheless, this spontaneous Ca_V_1.2 activation without action potential may be inefficient which is compensated by higher expression of Ca_V_1.2 channels to initiate the Ca^2+^ signaling for mechanotransduction. It is the case that we observed more robust Ca_V_1.2 expression in all tendons during early tendon growth before the formation of a stable myotendinous junction as well as in adult patellar tendons without myotendinous junction.

In summary, our data identified a novel role of Ca_V_1.2 voltage-dependent Ca^2+^ channel in nonexcitable tissue tendon and demonstrated that increased Ca^2+^ influx through Ca_V_1.2 promotes tendon formation predominantly by regulating tendon collagen fibrillogenesis. This was achieved through increased expression of the growth factor myostatin and a combination of differentially expressed fibril associated FACIT type XIV collagen, MMPs, TIMPs and other ECM proteins. These biochemical responses following increased Ca^2+^ signaling may cooperate in the adaptation of tendon in response to mechanical loading or tendon healing after tendon injuries. Pharmacologically, Ca_V_1.2 agonists, such as BayK-8644 or FPL 64176 can mimic the effect of Ca_V_1.2^TS^ to increased Ca^2+^ influx across the plasma membrane. Therefore, our data in this study highlights a potential therapeutic strategy to target Ca_V_1.2 channel and promote tendon formation and healing after injuries.

## Methods

### Mice

All animal studies were approved by the University of Rochester Committee for Animal Resources. *Ca_V_1.2^+/LacZ^* and *Ca_V_1.2^TS^* mice have been described previously(Chike Cao et al., 2017; Sergiu P Paşca et al., 2011; Kapil V Ramachandran et al., 2013). *Ca_V_1.2^+/lacZ^* mouse line carries a *lacZ* reporter with a nuclear localization signal under the promoter of *Cacna1c*, the gene encoding *Ca_V_1.2. Ca_V_1.2^TS^* mouse line carries a rat G406R TS-causing mutant Ca_V_1.2 cDNA which was knocked into the *Rosa26* locus with an upstream floxed stop codon to control the transgene expression by the *Cre-loxP* system. Homozygous floxed *Ca_V_1.2^TS^* mice were crossed with the transgenic *ScxCre* mice (provided by R. Schweitzer) to induce *Ca_V_1.2^TS^* expression and modulate the Ca^2+^ signals in tendon during development and growth. All tendon analyses were performed on mice at 1 month of age unless otherwise specified. Mutant mice or littermate controls of both male and female mice were analyzed unless otherwise specified.

### X-gal staining and histology

It has been descripted previously (Chike Cao et al., 2017). Briefly, visualization of *lacZ* expression was done by X-gal 5-bromo-4-chloro-3-indolyl-β-D-galactopyranoside) staining in whole mount embryos or limbs. For whole mount X-gal staining, embryos, forelimbs or hindlimbs were fixed in ice-cold fixation solution (2% paraformaldehyde and 0.5% glutaradehyde in 1× PBS) for 20 minutes (for embryos), 1 hour (for postnatal stage), or 2 hours (for adult stage), washed with 1x PBS for 3 times, each 10 minutes, processed with X-gal staining solution (5 mM potassium ferrocyanide, 5 mM potassium ferricyanide, 1 mg/ml X-gal, 2 mM MgCl2, 0.1% sodium deoxycholate, and 0.2% IGEPAL CA-630) in dark for 24 ~ 72 hours at 37°C. For histological analysis, whole mount X-gal-stained tissues were further decalcified with 14% EDTA at 4°C, 30% sucrose, and snap-frozen embedded with OCT compound (Sakura Finetek). Frozen sections (10 μm thickness) were counterstained with Nuclear Fast Red. For histology on fresh tendon tissue, tendons were isolated and immediately processed into 30% sucrose for 1 hour at room temperature, snap-frozen embedded with OCT compound. Samples were cross-sectioned at 10 μm thickness. Sections were air-dried for 1 hour at room temperature, fixed with 2% paraformaldehyde and 0.5% glutaradehyde in 1× PBS for 10 minutes, followed by Hematoxylin and Fast Green staining with standard protocols.

### RNA extraction and RT-qPCR

For total RNA isolated from tendon tissue, miRNeasy Mini Kit (Qiagen) was used. Achilles tendon and plantaris tendon were carefully dissected from control and *ScxCre;Ca_V_1.2^TS^* mutant mice, both sexes at 1 month of age. Tendons from each animal represented as one biological replicate without pooling tissues from different animals. Tendons were homogenized in QiAzol lysis reagent (Qiagen) using Biomasher II disposable micro tissue homogenizer and total RNA was purified following the kit instruction. Total RNA (500 ng) was reverse-transcribed to cDNA using cDNA reverse transcription kit (Applied Biosystems, Thermo Fisher Scientific) and qPCR with SYBR green Supermix (Bio-Rad). Relative expression was calculated using the 2^-ΔΔCt^ methods by first normalization to Gapdh (ΔCt) and second normalization to control samples (ΔΔCt). The primers used for tested genes were listed in Table S1, with *Mstn, Scx, Tnmd*, and *Gapdh* primers were previously described (Christopher L. Mendias et al., 2008).

### Collagen transmission electron microscopy (TEM)

Achilles tendons from control and *ScxCre;Ca_V_1.2^TS^* mutant mice at 1 month of age were used for TEM analysis. First, mouse hindlimbs were fixed in 1 x PBS containing 1.5% glutaraldehyde/1.5% formaldehyde (Electron Microscopy Sciences, Cat#: 15950), 0.05% tannic acid at 4 °C for overnight with gentle agitation. Achilles tendons were dissected out and post-fixed in 1% OsO4. After washing with 1 x PBS and dehydration in a graded series of ethanol, tendon samples were rinsed in propylene oxide, infiltrated in Spurrs epoxy and polymerized at 70 °C for overnight. Ultrathin sections at 80 nm were used for imaging using a FEI G20 TEM by the core service at MicroImaging Center, Shriners Hospital for Children, Portland. ImageJ was used for the measurement of collagen fibril CSA.

### Mechanical properties testing

Uniaxial displacement-controlled stretching at 1% strain per second until failure descripted previously (Korcari, Buckley, & Loiselle, 2022) was applied to Achilles tendons isolated from *ScxCre;Ca_V_1.2^TS^* mutant or littermate controls at 1 month of age for both sexes. Achilles tendon preparation followed the previous reported description with some modification (Sarver et al., 2017). Briefly, Achilles tendons (without plantaris tendons) were dissected out with one end attaching to the calcaneus bone and the other end with muscle. Tendons were wrapped in 1 x PBS-soaked kimwrap and stored at −20 °C until use. Prior to mechanical tests, tendons were thawed at room temperature, submerged in 1 x PBS, cleaned away of muscle to prevent slipping, placed with both ends between two layers of sandpaper, glued with cranoacrylate (Superglue, LOCTITE), and secured with a compression clamps. Each Achilles tendon was first quantified by its gauge length and CSA from 3 evenly spaced width and depth measurements from high-resolution digital photographs of both top and side views of the tendon (Olympus BX51, Olympus). Mechanical property testing was performed in a bath containing 1 x PBS at room temperature. A uniaxial displacement-controlled stretching of 1% strain per second was applied until failure occurred. Load and displacement were recorded, and the failure of each mechanically tested tendon was confirmed which often occurred at tendon mid-substance. Tendon peak load was taken as the maximum load prior tendon’s failure, while tendon stiffness was specified by the slope of the linear region from the load-displacement curve. Tendon tensile stress was defined as the recorded load divided by tendon CSA, while tendon tensile strain as the displacement divided by the gauge length. Tendon elastic modulus was calculated by the slope of the linear region from the plotted tendon tensile stress-strain curve. Tendon structural properties (stiffness, and peak load) and material properties (peak stress and elastic modulus) were determined from each Achilles tendon.

### Proteomics and data analysis

Mass spectrometry (MS) proteomic analysis was performed at the Mass Spectrometry Resource Laboratory, University of Rochester. Achilles tendons (combined with plantaris tendon) were isolated from *ScxCre;Ca_V_1.2^TS^* mutant or littermate control mouse at 1 month of age. Tendons from each animal represented as one biological replicate without pooling tissues from different animals. Trypsin (Thermo Scientific) was used to digest the tendon proteins followed by disulfide bond reduction with addition of 5 mM of Bond-Breaker TCEP solution (Thermo Scientific) and by alkylation of reduced cysteines with the addition of 10 mM of iodoacetamine. LC-MS/MS analysis was performed using a Q Exactive Plus mass spectrometer (Thermo Scientific). Raw MS data files were analyzed with PEAKS to identify protein composition. Searches were performed against the Uniprot mouse proteomes database (UP000000589). Search results were adjusted to 1% false discovery rate (FDR), filtering out peptides which had a p-value greater than 0.01. Z-scores were calculated from the normalized abundance of each protein to create heatmaps via GraphPad. Only proteins with significantly different abundance (p < 0.05) were used in the heatmaps. In addition, DAVID bioinformatics Resources (https://david.ncifcrf.gov/tools.jsp) was used for GO term enrichment analysis for proteins exhibiting 1.5-fold higher or 1.5-fold lower levels of expression in abundance and FDR *p*-value <0.05.

### Statistics

Statistical analyses were performed using GraphPad Prism 9.0 or OriginPro 8. Twotailed unpaired *t* tests were used to compare between mutant and control groups. Fold changes were calculated by dividing the value of the mutant group by the value of the control group. Increasing or decreasing changes were calculated by dividing the value of difference between the mutant group and the control group by the value of the control group and then multiplying 100. Kolmogorov-Smirnov test (http://www.physics.csbsju.edu/stats/KS-test.n.plot_form.html) was performed to determine if the size of collagen fibril differs significantly between groups. * *p* < 0.05. ** *p* < 0.01.

## Acknowledgments

We thank Douglas keene (MicrroImaging Center, Shriners Hospital for Children) for providing the TEM core service, Dr. Sina Ghaemmaghami (Mass Spectrometry Resource Lab, University of Rochester) for providing Mass spectrometry service and Dr. Edward Schwartz (Center for Musculoskeletal Research, University of Rochester) for his critical reading and helpful suggestions. This work was supported by NIH NIAMS R21AR075214 and P30AR69655.

---

**Table S1.**
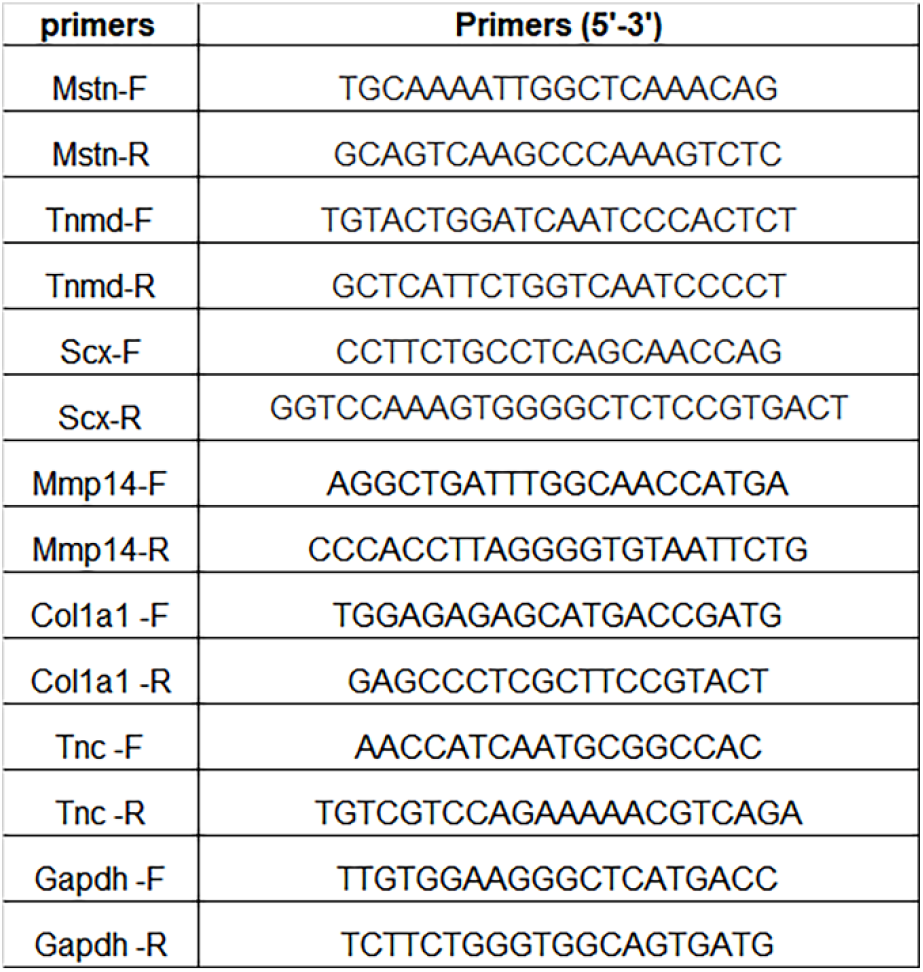
PCR primer sequences

## Notes

### Competing Interest Statement

The authors have declared no competing interest.

