## Supplementary for "Increased Ca^2+^ signaling through Ca_V_1.2 induces tendon hypertrophy with increased collagen fibrillogenesis and biomechanical properties"

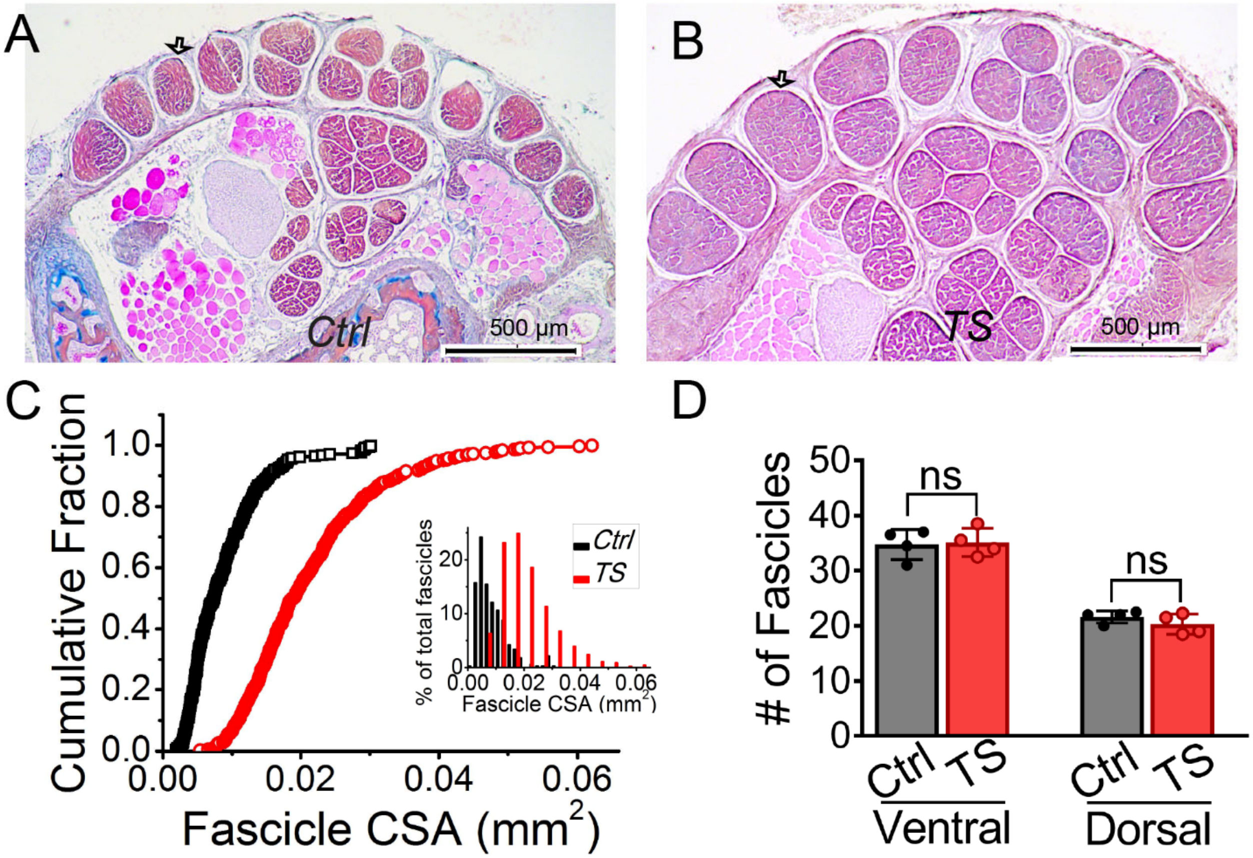


**Supplementary Figure. 1. *ScxCre;Ca$_V$1.2$^{TS}$* mice have affected tendon fascicle growth but not fascicle determination.** (A-B) Alcian blue Hematoxylin/Orange G staining of control and ***ScxCre;Ca$_V$1.2$^{TS}$*** mutant tail tendons. (C-D) Analysis of tail tendon fascicles by size (C) and number (D). Kolmogorov-Smirnov test shows that the tendon fascicle cross section area (CSA) in the *Ca$_V$1.2$^{TS}$* mutant is larger than in control mice, n = 4, $p < 0.001$, as also illustrated by the inset showing an altered size distribution.

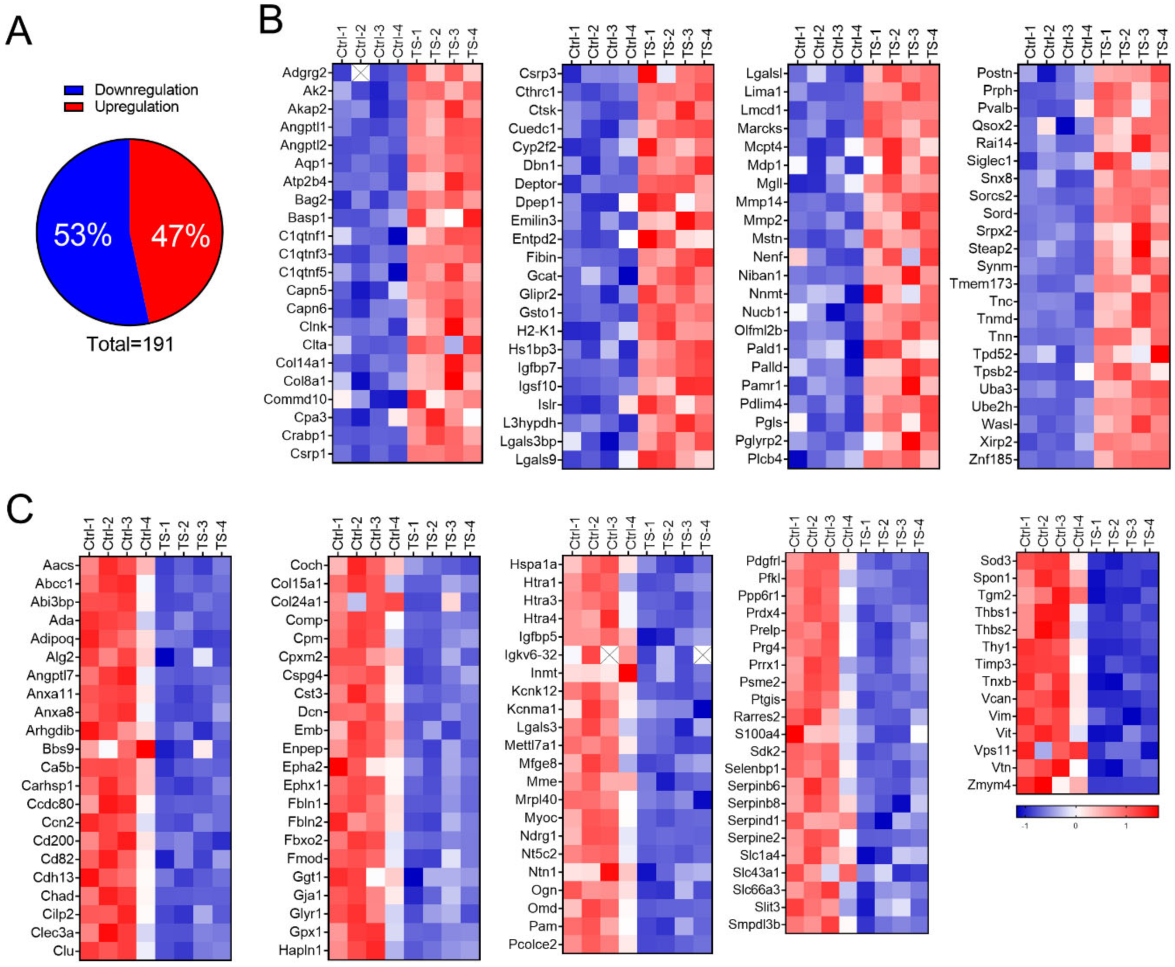


**Supplementary Figure 2. Upregulated and downregulated proteins in Achilles tendons in *ScxCre;Ca$_V$1.2$^{TS}$* mice.** Proteomic analysis was performed on Achilles tendon from control (*Cre$^-$;Ca$_V$1.2$^{TS}$*) and *ScxCre;Ca$_V$1.2$^{TS}$* mice at 1 month old. (A) Percentage of total proteins significantly altered in *ScxCre;Ca$_V$1.2$^{TS}$* mice versus control mice. (B) Heatmap showing all 89 significantly upregulated proteins in Achilles tendons of *ScxCre;Ca$_V$1.2$^{TS}$* mice versus control mice. n =4, Identification of proteins: > 1.5-fold change in abundance and FDR *p*-value <0.05 was considered significant. (C) Heatmap showing all 102 significantly downregulated proteins in Achilles tendons of *ScxCre;Ca$_V$1.2$^{TS}$* mice versus control mice. n = 4, Identification of proteins: < -1.5-fold change in abundance and FDR *p*-value <0.05 was considered significant.
